# Automated Analysis Pipeline For Extracting Saccade, Pupil, and Blink Parameters Using Video-Based Eye Tracking

**DOI:** 10.1101/2022.02.22.481518

**Authors:** Brian C. Coe, Jeff Huang, Donald C. Brien, Brian J. White, Rachel Yep, Douglas P. Munoz

**Author notes:** Author Contributions, BC, DB and DM conceived the study. BC created the analysis pipeline, wrote the paper, and created the figures. BC and DB created numerous analysis software applications in Matlab. JH defined post-processing of pupil area methods. RY contributed to the organization and verification of the dataset. JH, RY, and DM helped edit the manuscript. JH, DB, BW, RY, & DM were constant sources of feedback during creation of pipeline.

## Abstract

The tremendous increase of video-based eye tracking has made it possible to collect data from thousands of participants. Traditional manual detection and classification procedures for saccades and trial categorization is not viable for the large data sets being collected. Additionally, high-speed video-based eye trackers now allow for the novel analysis of pupil responses and blink behavior. Here we present a detailed description of our pipeline for collecting data, storing data, organizing participant codes, and cleaning data, which are fairly lab-specific but nonetheless important precursory steps in establishing standardized pipelines. More importantly, we also include descriptions of the automated detection and classification of saccades, blinks, ‘blincades’ (blinks occurring during saccades), and boomerang saccades (two saccades in opposite directions that occur almost simultaneously so that speed-based algorithms fail to split them), which are almost entirely task-agnostic and can be used on a wide variety of data. We additionally describe novel findings about post-saccadic oscillations, and provide a method to get more accurate estimates for end-points of saccades. Lastly, we describe the automated behavior classification for the Interleaved Pro- and Anti-Saccade Task (IPAST), a well-known task that probes voluntary and inhibitory control. This pipeline was evaluated using data collected from 592 human participants between 5 and 93 years of age, making it robust enough to handle large clinical patient data sets as well. In sum, this pipeline has been optimized to consistently handle large data sets obtained from diverse study cohorts (i.e., development, aging, clinical), and collected across multiple laboratory sites.

## Intro

Eye tracking has a long and rich history as a tool to study brain function (Leigh & Zee, 2015), including basic science as well as clinical studies. Video based eye tracking technology has advanced to a state that incorporates high frequency sampling and high spatial fidelity that is also minimally invasive. This has led to the opportunity to collect and integrate large datasets from multiple laboratory sites, thus new tools are required for automating and standardizing of data analysis and reporting. This pipeline needs to be centralized so that all data is analyzed in the same manner. In this way the well-known variability of user-based, hands-on detection and classification of events, like saccade onset, is replaced with an objective analysis and trial marking pipeline that is both reliable and repeatable.

Video-based eye tracking uses the pupil to estimate gaze position, and the pupil is defined by the ever-changing iris. This can detrimentally affect the spatial accuracy of video-based eye tracking because the orientation of the iris relative to the rest of the eye changes due to fluid dynamics and rotational acceleration. We describe a novel method to improve saccade endpoint accuracy. As many of the methods for these studies from our lab have been honed and repeated, it seems appropriate to provide a singular description for the techniques that have been built up over time and that we currently employ.

We have used highspeed (500Hz) video-based eye tracking to collect data from healthy participants across the life span as well as neurological and psychiatric clinical populations. Here we will only be addressing healthy controls to describe the analysis pipeline that automates and optimizes data analysis from video-based eye tracking. In this way we can extract the exact same measures of eye position, pupil size, blink probability, and task performance from hundreds of participants from numerous data collection sites.

## Methods

### Equipment

During the eye tracking, participants were seated in a dark room with their heads positioned comfortably in a head rest. Participants were seated so that their eyes were 60cm away from a 17-inch LCD monitor (1280×1024 pixels, 32-bit color, 60Hz refresh rate). An infrared video-based eye tracker (EyeLink 1000 Plus, SR Research Ltd, Ottawa, ON, Canada) was used to track monocular eye position at a sampling rate of 500Hz. When able, a 9-point array calibration and validation procedures was performed for each participant prior to the start of each task to ensure appropriate accuracy of eye tracking. In some situations where calibration was difficult, a 5-point array was used.

### Tasks

Although most of the collection, curation, & pre-processing described below can be implemented for multiple eye tracking tasks, here we concentrate on data from an Interleaved Pro- and Anti-Saccade Task (IPAST, see Figure 1) and some free viewing of common scenes (FreeView). The IPAST data is structured and gives specific behavioral metrics, like reaction time and error rate, whereas the FreeView data is unstructured and is conducive to recording a more robust variety of natural saccades. In general, these two tasks were collected on the same day. Each IPAST trial began with 1000ms of an inter trial interval (ITI) consisting of a blank black background (0.1cd/m2). After that, a central fixation point (FIX; 0.5° diameter dot, 44cd/m^2^) was displayed for 1000ms. The color of the FIX conveyed the instruction for each trial (green: pro-saccade trial, PRO; red: anti-saccade trial, ANTI). Following the FIX, the screen was again blank for 200ms (gap period, GAP). Finally, a peripheral stimulus (STIM; 0.5° diameter dot; gray, 62cd/m^2^) appeared 10° horizontally to the left or right of where the FIX was. On PRO trials, participants were instructed to make a saccade to the STIM as soon as it appeared (green arrow, bottom of Fig. 1). On ANTI trials, participants were instructed to look toward the opposite direction of the STIM (red arrow, bottom of Fig. 1). Trials where the first saccade after STIM onset was in the direction opposite of the trial instruction are known as Direction Errors (blue and orange arrows, bottom of Fig. 1). The saccade conditions (PRO or ANTI) as well as STIM locations (left or right) were pseudo-randomly interleaved with equal frequency. The IPAST experiment consisted of two blocks of 120 trials each with a short break in between, lasting approximately 20 minutes in total. Special codes were logged in the eye tracker’s raw data file (*.EDF) to represent the timing of the events described above and the nature of each trial, as well as screen metrics such as screen size and distance between the participant and the screen. The timing of all stimuli was verified by external photosensor.

**Figure 1:**
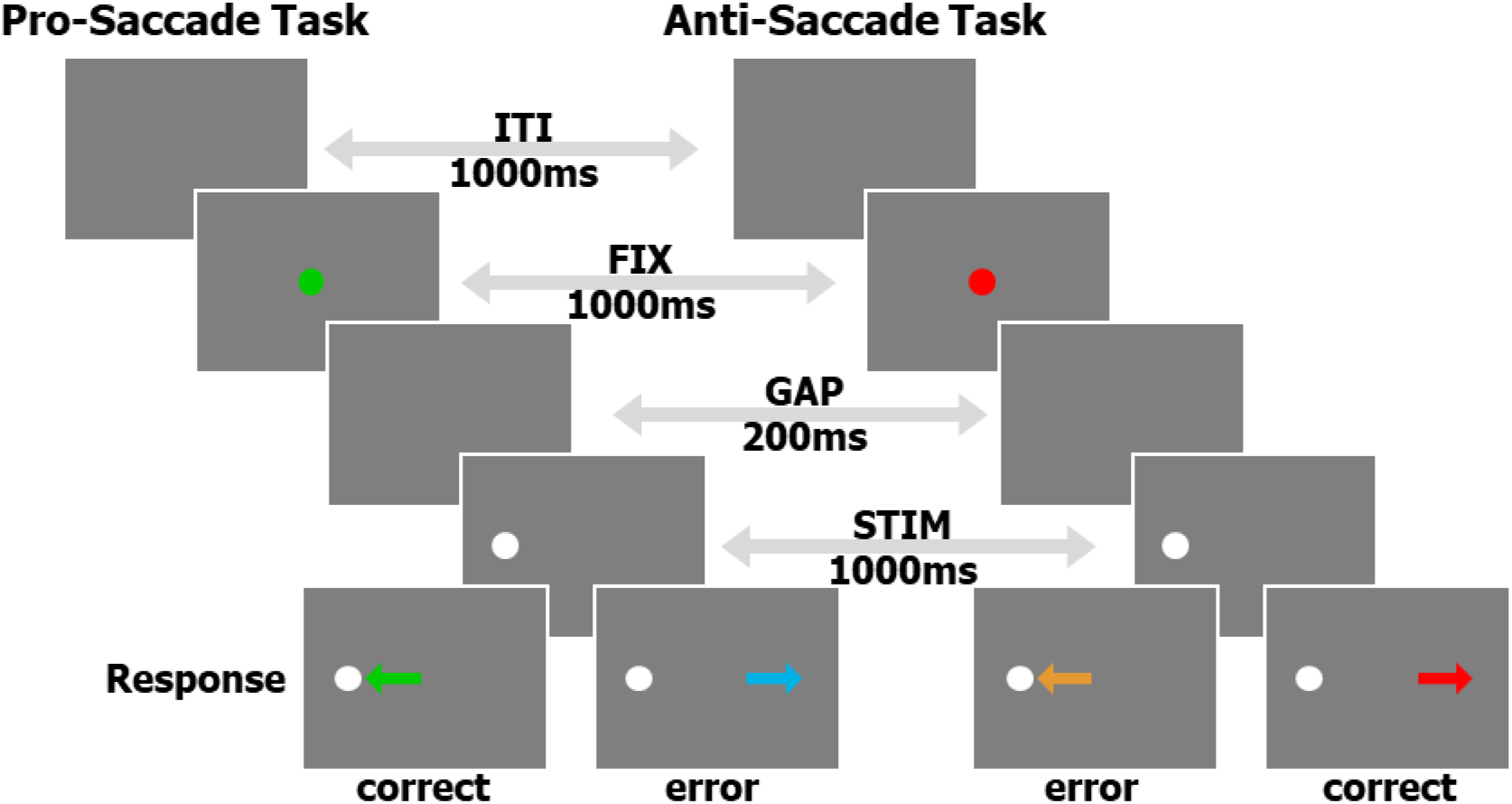
A schematic of the timing and visual aspects of the IPAST.

For FreeView data collection, videos were displayed on the same monitor as the IPAST using custom software in Ubuntu 13 that could interface with the eye tracker via the SR Research API. All participants viewed a total of 10 movies at 30fps and with no audio. Each movie was contained in an independent file, was approximately 1 minute in duration, and consisted of 15~17 video clips that were 2~5 seconds each in duration. The video clips featured scenes that may contain humans, animals, buildings, cars, and/or natural scenes. The clips in each movie instantaneously changed from one clip to the next, and were presented in a fixed sequence within each movie, but the order of the first 5 movies and then the second 5 movies was randomized between participants. The task required no instructions, and the participants simply viewed the movies. The timing of all stimuli was verified by external photosensor.

### Participants

All experimental procedures have been assessed and accepted by the Queen’s University Health Sciences and Affiliated Teaching Hospitals Research Ethics Board. Individuals between the ages of 5-93 were recruited from the greater Kingston, Ontario, area using online and newspaper announcements. The participants reported no history of neurological or psychiatric illness and had normal or corrected-to-normal vision. An informed consent form was obtained from all individuals over the age of 18. An assent form, in addition to a parent’s or guardian’s informed consent form, was obtained from all individuals under the age of 18. Study sessions took approximately one hour. For their participation, participants were remunerated $20 CAD.

In total, 592 participants across a dynamic age range contributed to the current dataset (see Fig. 2). All participants performed at least one block of the IPAST and had at least 30 viable saccade trials from both the PRO and the ANTI tasks. Viable saccades are described in the Saccade Classification section below. 407 participants also completed the FreeView task.

**Figure 2:**
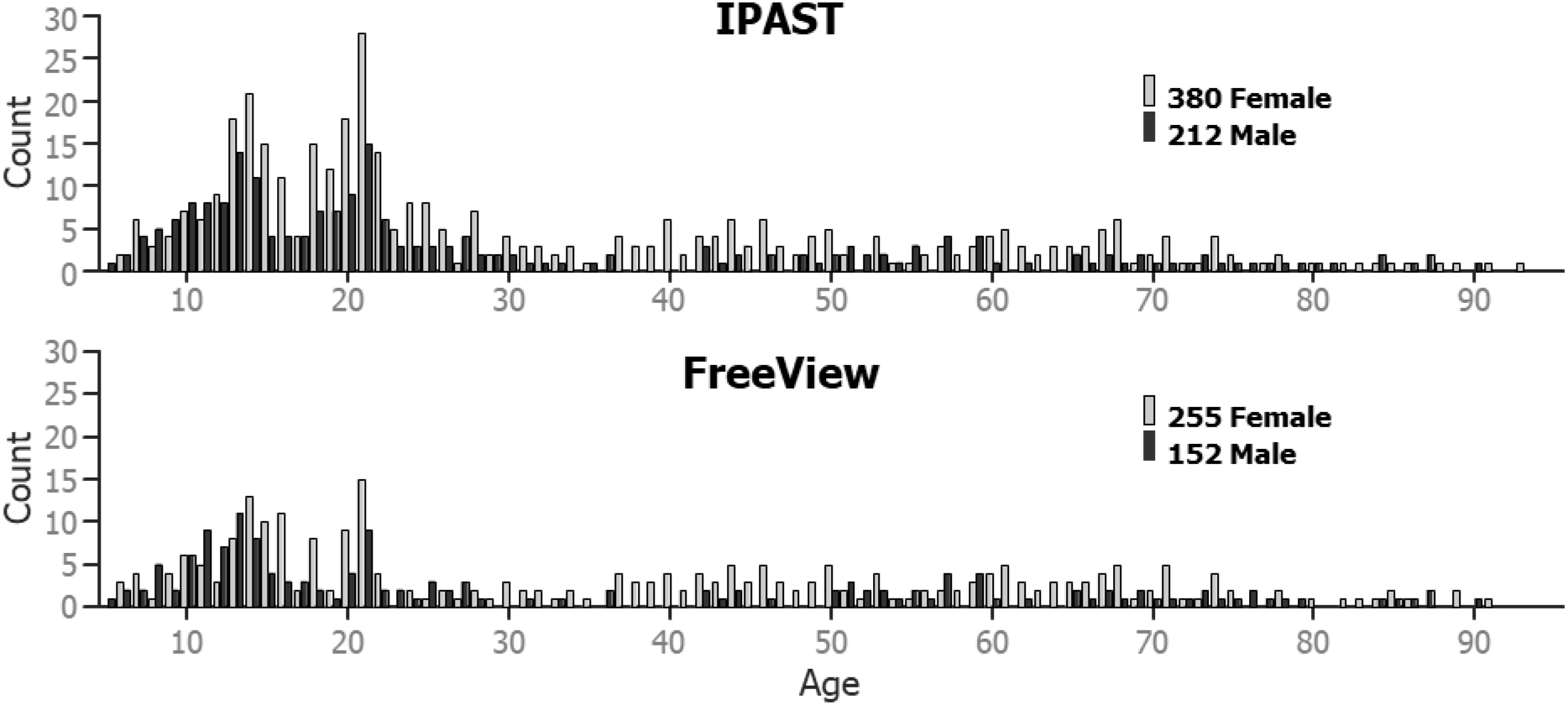
Age based histograms of the participant data base, per gender and task.

### Collection, Curation, & Pre-Processing

All raw eye tracking data files (*.EDF) were stored in task specific folders (i.e., each run/block had its own folder) inside of one participant folder, which was labeled using a 4-digit participant identifier and the date of recording (yyyymmdd, in descending order for easier sorting). An online database of available participant identifiers was setup so that numerous data collection sites could work in unison to assure that each participant was assigned a unique identifier.

#### Collection

Basic demographic information about each participant was digitally collected immediately prior to each of the two IPAST runs/blocks via experimenter input to a custom front-end program that was written in Java. This program performed error checking on our naming conventions, launched the IPAST automatically, and then saved the eye tracking data (*.EDF file) and the demographic information (*.txt file) in the previously described task specific folder to reduce errors in data collection and participant identification. FreeView data collection took place following the IPAST and was stored in its task specific folder and all three task specific folders (2 IPAST and 1 FreeView) were stored inside one participant folder.

#### File Conversion

All data processing was completed using custom software in MATLAB (The Mathworks, www.mathworks.com). Raw *.EDF files were converted to MATLAB files via custom code using EyeLink API. This was done in a task-agnostic fashion thus the process can be used on any EDF file. There were minimal changes to the data during this conversion. The first timestamp in each file was stored separately and then subtracted from all other timestamps in the file. This allowed for the data size to be reduced to a ‘single’ (32-bit precision) instead of a ‘double’ (64-bit precision). Other scores were similarly reduced in size while still maintaining the full range of values. These small changes made the converted data size approximately 25% of the original data size and drastically decreased the time it took to load and analyze the files. Timestamps, horizontal and vertical eye position, and pupil area data (T, X, Y, A, respectively) were collected at 500Hz. All task events were recorded with a corresponding timestamp in the *.EDF file. All event timestamps were rounded to match the recording rate (i.e., multiples of 2) for easier alignment with the eye data. Raw X, Y, and A data were recorded in pixels. The converted data file (*_EDF.mat) was then saved in the original raw data folder for quick and easy loading.

#### Cleaning & Combining

The EyeLink 1000 pauses recording in between trials, so the data is not continuous throughout the entire recording, thus further analysis is done on a trial-by-trial fashion. The next step was to load the converted *_EDF.mat file using custom task-specific functions that would first separate the T, X, Y, A data into individual trials, using common event stamps implemented by the EyeLink Experiment Builder task creation software. Although extremely rare, timestamp gaps or repeats were fixed if they occurred; either by linear interpolation at 2ms for the missing timestamps, and the corresponding X, Y, and A, or by removing the duplicate timestamps and the corresponding values, respectively. The converted data file (*_EDF.mat) for each IPAST block was combined with its paired metadata file (*.txt) created by the front-end Java program. For each FreeView block we used a metadata file from an accompanying IPAST block in the same participant folder as they were run on the same day. The pixel-based X and Y data were transformed into degrees of visual angle using screen size and the distance from the participant to the screen that we digitally added to the raw *.EDF file. The EyeLink 1000 has the ability to update the initial calibration at regular intervals between trials during the experiment, known as a ‘drift-correct’. For each of the intervals between ‘drift-corrections’ we determined the mode of eye position during a known fixation period for each trial to best asses the participants actual fixation (Harris et al.,1990). This was used to create a post-hoc X & Y adjustment to better estimate the actual fixation of each participant. This new data structure was then entered into an automated pipeline that performs pre-processing (described below) and then task-specific trial-by-trial auto-marking for behavioral markers (described in the next section).

Smoothing is a common method to help clarify high frequency data. We chose to use MATLAB’s *filtfilt* function that “performs zero-phase digital filtering by processing the input data, x, in both the forward and reverse directions.” (MATLAB help documentation). For a review of different smoothing applications on different recording rates see Mack et al. (2017). We used a box shaped transfer function coefficients for the *filtfilt* function, which is similar to a moving average with a given width. Thus, the larger the width, the larger the window, and the greater the smoothing. This was used for all filtering as this was the simplest form, treated all data points equally (saccades or otherwise), had the fewest assumptions, and had the smallest effect on the starts and ends of changes in velocities (see Mack et al. (2017) Fig 3). The MATLAB code was made to be dynamic by adjusting a single variable, the width:

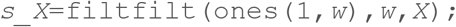

where *w* is the width of the window for smoothing, *X* is the data to be smoothed, and the ‘`ones`’ function giving equal weight to all data points in the window.

**Figure 3:**
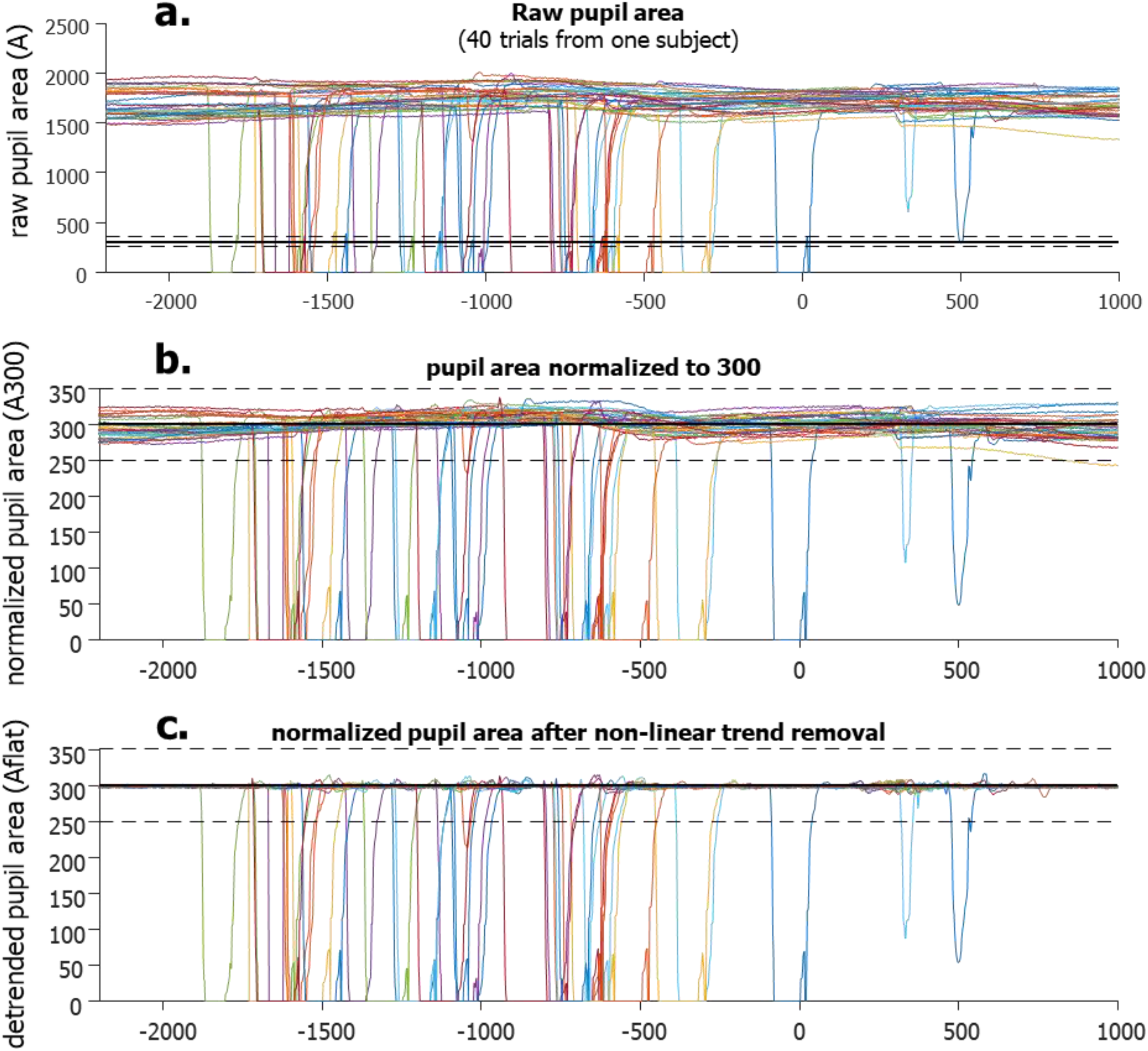
Blink detection Algorithm. Panel A shows raw data (in pixels) from 40 trials of one participant. Panel B shows data that has been normalized (trial by trial) to have a mean of 300 (now in arbitrary units). Notice that in some cases the natural fluctuations make using a fixed threshold (dashed horizontal line) for blink onset untrustworthy. e.g., the far right end of the yellow trace dips below our 250 limit. If we were to lower the limit, small eye loses (orange trace just prior to −1000ms) would be missed. Panel C shows the benefits of non-linear trend removal, clearing the way for a fixed threshold for blink detection. It can include partial blinks and not be triggered by large fluctuations, as they have been removed.

In a similar manner, differential calculations for instantaneous X and Y velocities were also done in a zero-phase manner. To calculate an instantaneous differential for a given point in time, the previous data point was subtracting from the following data point and then divided by 2. For the first and last data points, a simple 2-point difference was used.

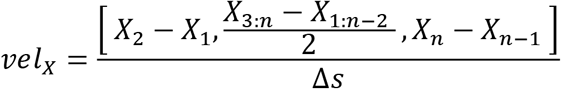

Where X is the vector of horizontal position (replace with Y for vertical position), *n* is the number of data points, and Δ*s* is the difference in time in seconds between each datapoint (Δ*s*=0.002 for 500Hz recording). Eye speed in degrees per second was determined via the usual Euclidean process using both of the 3-point smoothed eye velocity vectors.

#### Pre-processing

Once the data was separated by trials using custom task-specific functions, the data was run through the following two pre-processing steps, which are task-agnostic.

##### Blink detection & definition

While video-based eye tracking is by far the easiest and least invasive form of eye tracking, it suffers from data loss during several situations. Blinks can cause full, or even partial, occlusion of the pupil, which results in data loss. At first a nuisance, this data loss has recently become useful for studying a vast variety of blink behaviors. This data loss during eye tracking can occur for a number of different reasons, not just blinks. For example, eye makeup like mascara can fool the pupil detection algorithm into including the lashes into the pupil estimate, creating an unstable and noisy signal. Thus, the most important issue is not to detect this loss but to classify it properly as either a true blink or some other loss.

In order to better isolate when data lost was due specifically to blinks, several steps were taken to clarify the pupil area data. As the pupil area data (A) can be quite variable (Fig. 3A) the first step was to normalize each trial to have the same average area.

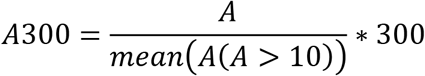

The raw pupil area values were the number of pixels of the eye’s image in the tracking camera, and generally ranged from the hundreds to the thousands. Thus, values below 10 were not included in the definition of the mean and the number 300 was arbitrarily chosen. This made the non-zero mean of the pupil area fixed to 300 for each trial (A300, Fig. 3B). The next step was to model the low-frequency modulation of A300 over time so that it could be removed to create a flattened A300. The goal of this step was to help clarify the sections of data that were indicative of eye loss. We smoothed the velocity (i.e., change in A300) using a 3-point kernel, and this smoothed velocity (sVEL) was used to flag high velocity data for removal and replacement. A model of the low-frequency modulation was created using a copy of the A300 data where the high velocity (sVEL>1000) and an abnormal area (A300<200 or A300>400) data were replaced using liner interpolation between the preceding and following data points. This model was then smoothed using a large 50-point kernel to help remove the high-frequency modulation of the area signal. This model was subtracted from the A300 to remove low-frequency, non-linear trends to create a flattened profile that exaggerated high velocity changes in area (Aflat, Fig 3C), which greatly helped in detecting the start and end points of lost data. These were defined as Aflat<250 or Aflat<350 (black dashed lines in Fig 2). We noticed that data loss was preceded by, and followed by, drastic fluctuations in pupil area, indicating that an actual blink lasted longer than the loss of data. Typically, the A300 would drop off quickly just prior to data loss. This was presumably due to the eyelid progressively occluding the pupil, thus reducing the number of pixels the eye tracker delineated as the pupil. Likewise, after the data loss, the A300 would recover from near zero and waver around 300, as more of the pupil became visible to the camera again. To better define the actual duration of a blink, these fluctuations were considered part of the blink. Once start and end points of the data loss were detected we used the smoothed absolute velocity (saVEL) of the A300 to determine the start and end of the full blink using a dynamic threshold per trial, similar to using speed for saccade detection.

We noticed that blinks generally have stereotypical durations and patterns on the pupil area data. We also noticed that each individual’s blinks affect the data slightly differently. We used the duration of data loss to categorize whether a loss in data was due to a blink or due to some other interference. Although the blink detection analysis is task-agnostic, task specific software was used to visualize the output. For example, figure 4 shows a screenshot of custom software used to visualize an individual’s blink behavior during the IPAST. Trials are reordered by task (ANTI over PRO) and by location of the STIM (left over right). The trials start with the ITI, followed by the colored FIX. Blinks are illustrated by black and grey lines indicating the duration of the data loss and the full blink, respectively. Task related saccades are indicated between 90ms and 800ms after STIM onset and are color coded to match Figure 1. For this subject, and many others, the ITI and FIX intervals show an increased number of blinks in an organized manner, despite no such instructions to participants. The main window also shows blink probability (not just onset) histograms over-laid at the bottom. The panel on the lower left shows histograms of blink durations and are color coded by task. This individual was chosen for their consistent blink behavior, which was common among most participants. However, there were some individuals who rarely blinked at all.

**Figure 4:**
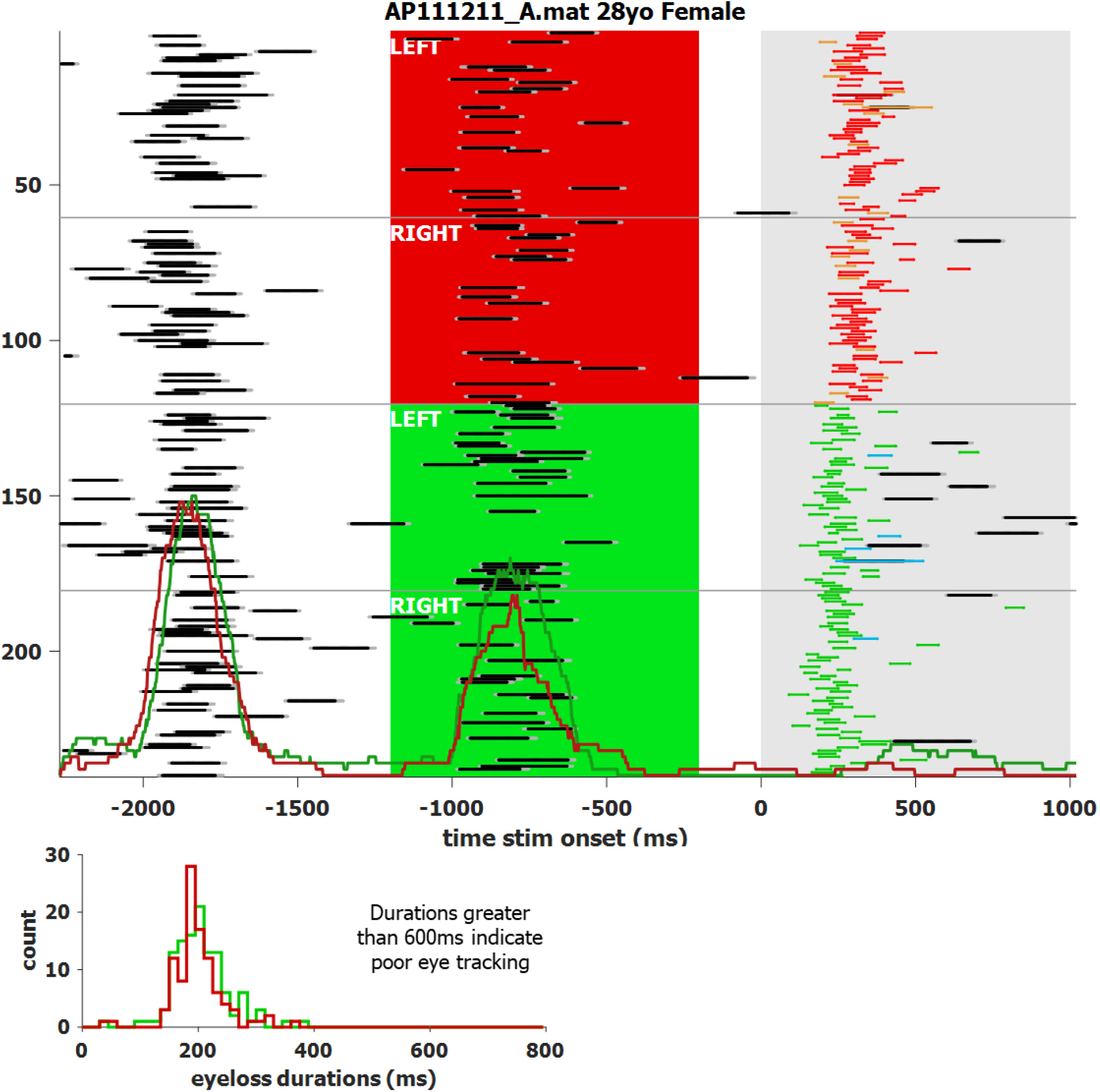
A screen shot of a custom application to visualize the blink behavior of a single participant with a high blink rate. Data loss is indicated by black lines and bordered by grey to indicate full blink length. Task related saccades, between 90 and 800ms, are color coded to match figure 1. Red and green histograms overlaid in main window indicate probability, not just onset, of a blink for anti and pro saccades, respectively. Their heights are proportional to number of total trials. Trials are re-ordered by task (anti over pro) and STIM location (left over right).

##### Saccade detection & definition

Given relatively clean eye tracking data, the detection of the onset of a saccade is a straightforward process of determining a speed threshold and then finding the first data point that is above it, with consecutive data points remaining above it for a given duration. The background noise during a fixation epoch, where eye speed was below a fixed threshold of 50°/s, was used to determine the dynamic threshold for each trial. Thus, noisier trials had a higher threshold to avoid numerous false positives. We used data below this fixed threshold to find the mean and standard deviation of the background noise. Our dynamic threshold for each trial was defined as the mean plus 2.5 times the standard deviation, but never less than 20°/s. The eye speed had to remain above this dynamic threshold for 10ms.

The end point of saccades, however, can be quite difficult to properly determine due to well-known aspects of video-based recording (Hooge et al., 2015; Mardanbegi et al., 2018; Nyström et al., 2013) and how they differ from the classical search coil technique (Frens & van der Geest, 2002; Kimmel et al., 2012).

Video-based eye tracking uses the pupil to estimate gaze, and the pupil is defined by the everchanging iris. The iris not only expands and contracts to change the diameter of the pupil, but also the orientation of the iris relative to the rest of the eye also changes due to fluid dynamics and rotational acceleration. This means that the pupil/iris orientation does not always lineup with the orientation of the eye and gaze. The forces on the iris have been described in detail before (Bouzat et al., 2018; Jacobi & Jagger, 1981). To summarize briefly, the lens and iris structures are situated in between two independent fluid bodies: the aqueous humor in front (light blue, Fig. 5), and the much larger vitreous humor (light grey, Fig. 5) that fills most of the eye, and these two systems do not connect. The vitreous humor is a closed (jelly-like) system that makes up the center-of-mass for the eye, whereas the aqueous humor is a flowing system (more like cerebrospinal fluid) and is on the periphery of the center-of-mass.

**Figure 5:**
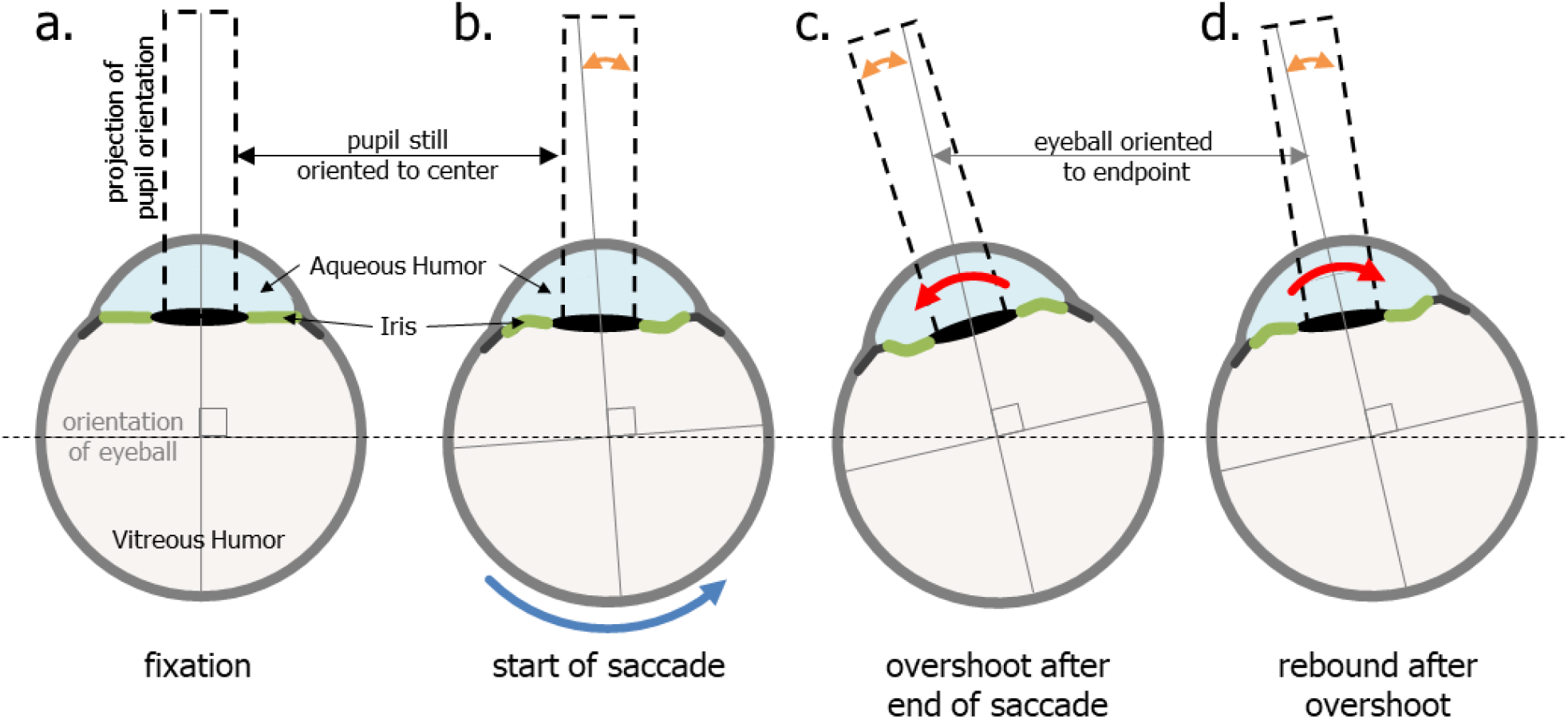
Shows the effect of rotational acceleration and deceleration on the shape of the iris and the orientation of the pupil (dashed black rectangles indicate orientation) in relation to the rest of the eye. The orange double headed arrows in panels b, c, d show the discrepancy between the orientation of the eye (which indicates actual gaze) and the orientation of the pupil (used by video-based eye trackers to estimate gaze). Panel B illustrates how the effects of inertia of the Aqueous Humor may cause a delay in the motion of the iris/pupil, though thought to be minimal. The last two stages are Overshoot(C) and Rebound (D) that alternate and diminish with each cycle until equilibrium (seen in fixation) is once again reached. This sloshing of the Aqueous Humor can cause, or increase, the well know post-saccadic oscillations previously reported by several papers. (Partially inspired by Jacobi & Jagger, 1981)

This arrangement causes the flexible iris to create something the field of Fluid Dynamics refers to as a Fluid-Structure Interaction. The rotational acceleration and deceleration of the eye creates a Slosh Dynamics Problem^1^ between two bodies of fluid with a flexible structure in between. A full and proper discussion of fluid dynamics is beyond the scope of this paper; however, the mere acknowledgement of these issues enables a better understanding of the mechanisms of the iris when the eye moves. Because the iris is flexible, the rotational acceleration of the eye and the inertia of the aqueous humor induces pressure strain on the solid structures of the eye that lie between the two liquid parts of the eye. Upon reaching a fixed (or zero) speed, this pressure is oscillatory and diminishes with each cycle until equilibrium is once again reached^2^. The clearest visualization of this oscillatory motion of the iris is seen in the condition known as Iridodonesis^3^ (numerous videos can be found on YouTube). Importantly, even healthy eyes have this oscillatory motion, albeit to a much lesser degree, and it interferes with the proper detection of the end of an eye movement.

> *To better illustrate the effects of the post-saccadic slosh, data from the FreeView task was used, as there was a larger variety of saccades during this task as compared to the IPAST. Figure 6 shows saccade data from 2 individuals from our data set; one with the most slosh detected (74yo Female) and the other with the least (46yo Female). The top two panels show a Fixed Saccade Map (FSM) where each saccade starting point was plotted from the center. The bottom two panels show speed traces of only rightward saccades of approximately 10°, in order to make cleaner comparisons. If only a fixed threshold was used to detect the end of the saccade, in the same way as detecting the start of the saccade, only the blue data points would have been considered part of the main saccade. The following supra-threshold data points would have been inappropriately classified as micro saccades. The red points indicate data that was added to the main saccade by detecting slosh generated post-saccadic oscillations (PSO) that were supra-threshold (e.g., >20°/s), close in time to the main saccade (<40ms between), and smaller in amplitude (between 0.5 and 5°). Ignoring these oscillations generally led to inaccurate saccade end points. These were often hypermetric, as the eye would stop but the iris would overshoot due to the fluid dynamics discussed above. For many people, including these PSO increases the saccadic end-point accuracy, but it clearly overestimates the duration, as seen in the bottom two panels of Figure 6. Of particular interest is that the horizontal saccades have a predominantly horizontal wobble whereas oblique saccades oscillate in a circular pattern, indicative of independent vertical and horizontal slosh patterns.*

**Figure 6:**
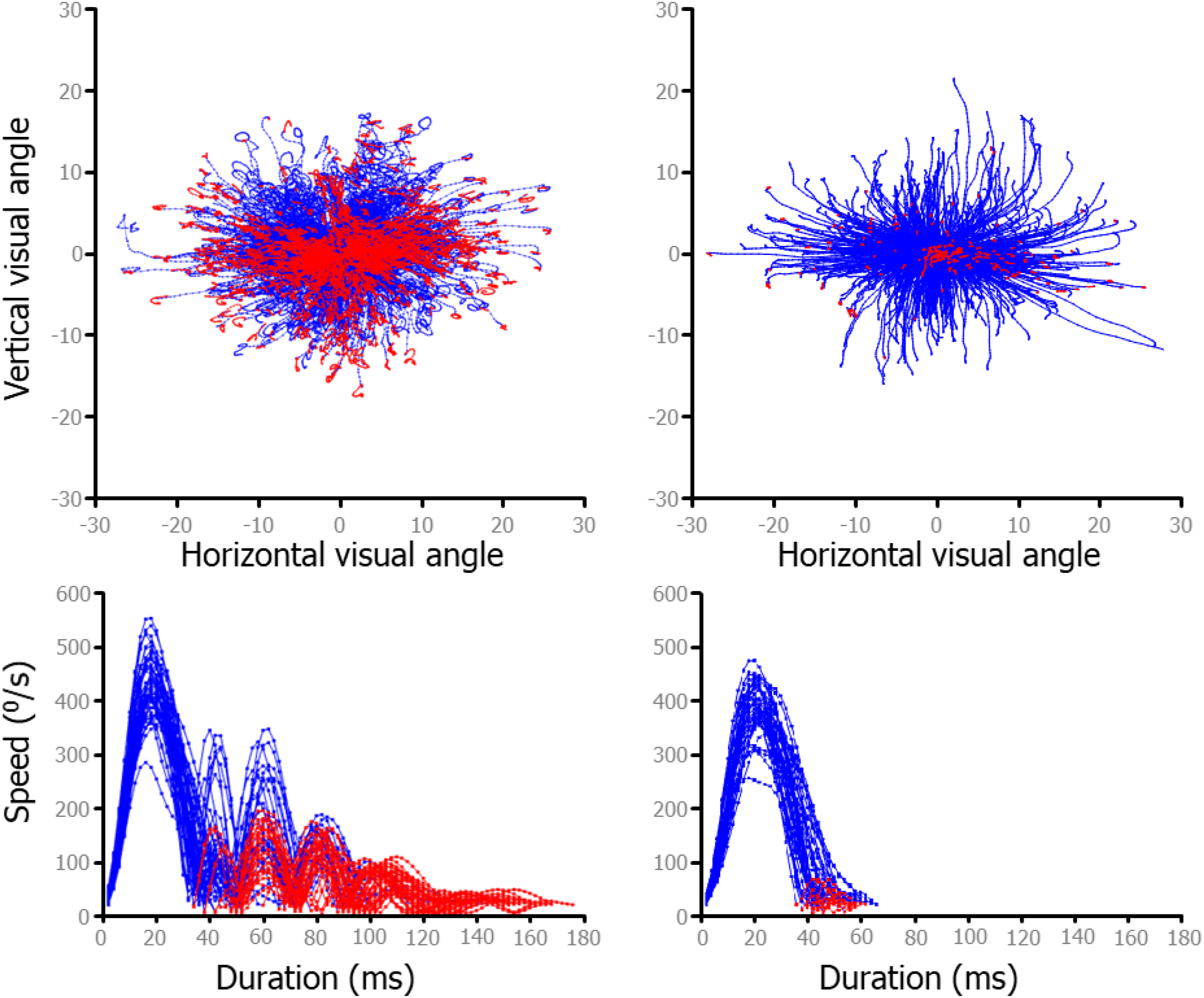
Fixed Saccade Mapping (FSM) and speed plots from free-viewing data. Red data indicates the part of the saccade after the speed dropped below threshold, but due to post saccadic slosh, motion continued. Some individuals have much more mobility of the iris, leading to greater post-saccadic slosh. Panel A shows a great deal of slosh and Panel B shows very little post-saccadic slosh. Panels C&D shows speed plots for a sample of each individual’s (directly above) saccades. To make sure to compare similar saccades, only rightward saccades made within ±22.5^0^ of 0^0^ and with amplitudes from 8 to 12 degrees were plotted.

##### Saccade table

For each block, of either task, a table of saccade information was created. Each row was one saccade with columns indicating: trial, start and end point, peak velocity, acceleration, amplitude, angle, and duration. Additional information was also included and is discussed below.

For the vast majority of saccades, the onset and offset (with the understanding of the post-saccadic slosh) are relatively easy to detect and define. However, two prominent exceptions still exist, one of them being more common in the IPAST. During an ANTI trial, participants occasionally made what we call ‘boomerang’ saccades. These are eye movements where the initial trajectory and the final trajectory are diametrically opposed and are indicative of simultaneous planning of two saccades (McSorley et al., 2016). This happens most frequently when a saccade is triggered to the error location but is instantaneously corrected and redirected back to the correct location, or sometimes returns to the fixation point and is followed by another saccade to the correct direction. With these boomerang saccades the velocity never drops below the speed threshold and thus automated algorithms would define this action as one single saccade, which would start at the fixation and end at the correct location. Or, in the case where the boomerang saccade returned to fixation, as a micro saccade that starts and ends near fixation followed by a correct saccade. Thus, this would lead to an incorrect trial being marked as a correct trial (see IPAST Auto-Marking). After the start and end of each saccade is detected, another step is used to detect if the initial direction and final direction are opposite and align with error and correct locations. This singular saccade is then split at a mid-point where speed is at its lowest. Doing so revealed both error-to-correct boomerang saccades and correct-to-error boomerang saccades, the latter being much more rare.

There were also incidents where eye tracking was lost during an eye movement. This could be due to a blink or due to other eye tracking issues. These ‘blincades’ were included in the table of saccades with a tag indicating that some data was lost during the change in eye position.

In some situations, a normal blink can lead the eye tracker to falsely indicate an upward ‘blincade’ with data loss, followed by a downward ‘saccade’ as the tracker regains the pupil. This would lead to mis-categorization of a blink where no movement took place as two saccades: a blincade upward immediately followed by a noisy saccade back down to the original position. For individuals who blink more frequently, this resulted in numerous fixation-break false positives (see IPAST Auto-Marking), and highly aberrant saccade metrics. In the situation where the automated process found an upward ‘blincade’ immediately followed by downward ‘saccade’ and the end point was within 2° of the start point, these two events were combined and marked as a blink, not two saccades. If the end and start point were farther apart than 2°, the two events were combined as a single ‘blincade’ with a linearly-interpolated trajectory from start to end.

As video-based eye tracking can be noisy, we know that some of the saccades are misidentified, thus a special process was performed to indicate how consistent the saccades were for each participant. For each block of data, Main Sequence (peak velocity vs amplitude and peak velocity vs duration) was calculated and modeled using a smoothing spline in MATLAB.

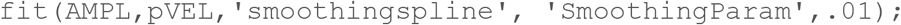

Saccades with known issues like missing data, boomerang saccades, or blincades, were removed from the fitting process. Then Z-scores were calculated for each residual. This fit was used to calculate the Main Sequence Z-scores (MaSeZs) that were added to the saccade table discussed above. These MaSeZs can be used to rank how ‘normal’ a saccade was for that participant and whether it’s metrics should be used to calculate eye movement scores. For example, when calculating a participant’s mean peak velocity only saccades with a MaSeZs less then 3.29 (p<0.001) was used, as a Z-score greater that that is indicative of some eye tracking issue where the behaviour can be detected but the instantaneous eye tracking was fouled.

### IPAST Pipeline

Up until this point the pre-processing steps described could be done with most tasks with minimal task-dependent alterations (e.g., identifying trial start and end points for each trial, and the timing of fixation epochs). From this point on, however, everything is task, and perhaps lab, specific so subsequent analysis descriptions will be more about theory and objective than code or formula oriented.

#### Pupillometry

In recent years there has been an increased interest in pupil size, as it is readily available with video-based eye tracking. Unlike the task-agnostic methods for blink detection, for the IPAST blink analysis the raw pupil area data, in pixel count, is used. Pupil size is a sensitive measure and can be affected by lighting conditions, blinks, noisy data, and eye position. To avoid distortion of pupil measures due to movement and position, pupil size analysis was done during the FIX and GAP periods for each trial and with several limitations. Only trials when the eyes were stationary (no saccades over 2°), looking in a central location (on or near the FIX), and without too much data loss (<200ms in duration) were used.

To capture the dynamics of the pupil response, we calculated pupil measurements for baseline, constriction, and dilation components (Fig. 7) for each trial. In response to the FIX onset, there was a standard pupil response that began with constriction and was followed by dilation, prior to the onset of the STIM. Baseline pupil size was calculated by averaging the pupil size during the epoch of 150 to 200ms after FIX onset, which is prior to any pupil response. Pupil response onset latency was defined as the earliest point at which pupil velocity significantly differed from the baseline, calculated using a 20ms sliding window. The Constriction Amount was calculated as the difference between baseline pupil size and the pupil size at maximum constriction, and the timing of this nadir in pupil size was defined as the maximum constriction time. The Dilation Amount was calculated as the difference between pupil size at maximum constriction and the mean pupil size just prior to STIM onset (−50 to 0ms). Pupil dilation velocity was calculated as the mean pupil velocity just prior to STIM onset (−50 to 0ms).

**Figure 7:**
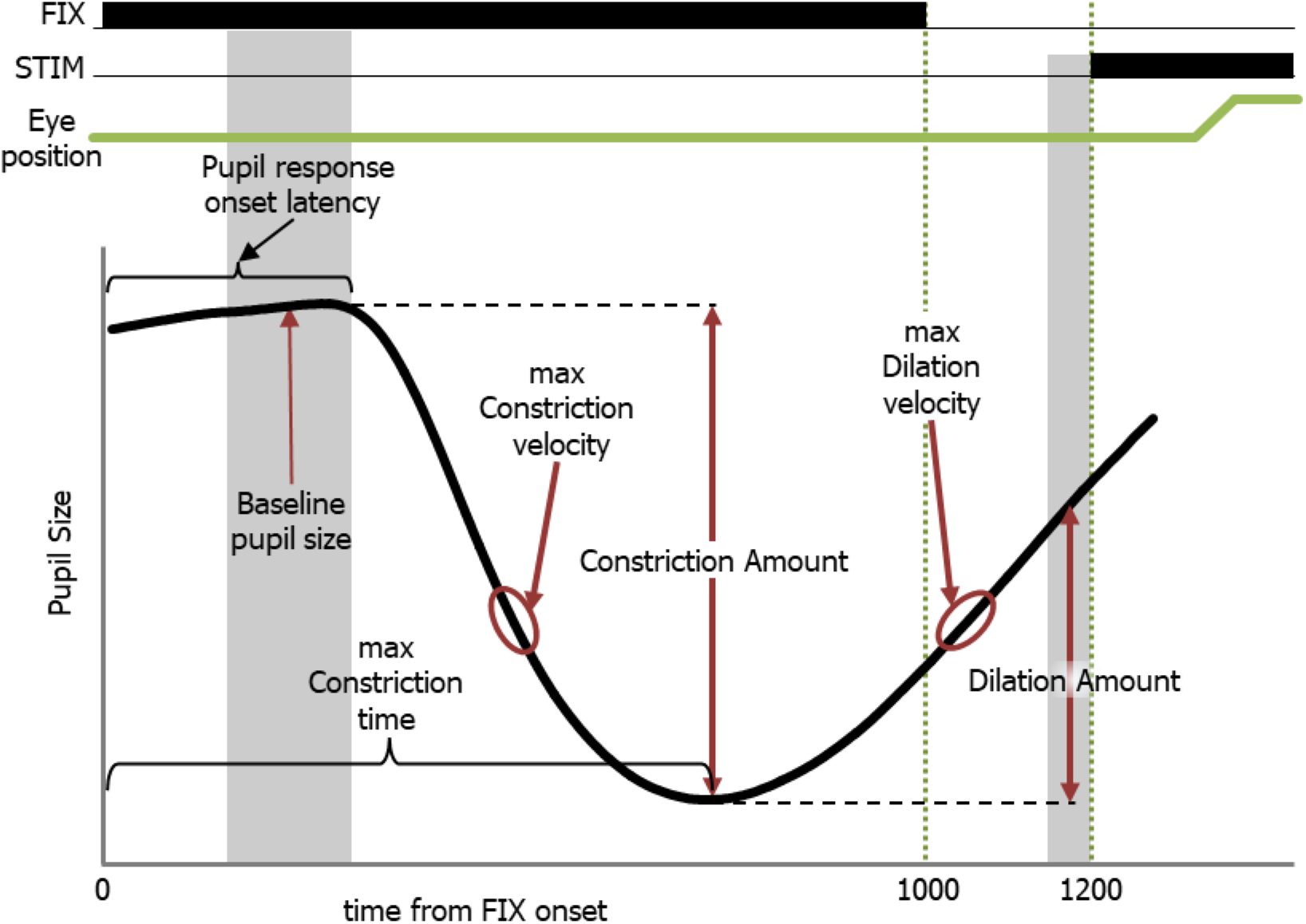
A sematic of the definitions for pupulometric measures during the IPAST.

#### IPAST Saccade Classification

All detected saccades were classified by when they occurred and their start and end positions. For viable saccades (see Table 1 & Fig 8), Saccadic Reaction Time (SRT) was defined as the time between the appearance of the STIM and the onset of the first subsequent saccade. Saccades towards the two potential stimulus locations that occurred during the GAP period were defined as anticipatory because they were equally likely to be either correct or direction errors, suggestive of a guessing behavior (Munoz et al., 1998). We determined that the timing of the earliest possible visually-triggered saccade was 90ms after STIM appearance. Prior to this time in the PRO task, correct responses and direction errors are equally present, but after this time correct responses are triggered in excess of 95% of trials. Thus, the anticipatory window is from −110 to 89ms (the GAP period but shifted by 90ms, light-green shaded area in Fig. 8). Express and Regular latency saccades have been defined and discussed previously (Fischer & Boch, 1983; Fischer et al., 1984) and are separated for reasons discussed in Coe & Munoz (2017). Here we delineate between the two latencies at 140ms. Saccades that take place after 800ms are extremely rare but do occasionally occur. These extremely long SRT responses were used to classify the trial-type (see below), but as they represented the participants inattention, they were removed from measurements of latency and saccade metrics.

**Table 1.** shows the trial categorization performance of the IPAST pipeline on data from 592 participants who attempted 137,896 trials.

| **Trial Type** | **Trial Description** | **Count** | **Percentage** |
| --- | --- | --- | --- |
| Not marked | trial has not been marked | 55 | 0.04% |
| Correct Pro-Saccade | A correct Pro-saccade between 90ms and 1000ms after STIM appearance | 58161 | 42.18% |
| Correct Anti-Saccade | A correct Anti-saccade between 90ms and 1000ms after STIM appearance | 46010 | 33.37% |
| Pro-saccade Direction error | An erroneous saccade to the Anti-location during the Pro-saccade task between 90ms and 1000ms after STIM appearance | 1612 | 1.17% |
| Anti-saccade Direction error | An erroneous saccade to the STIM during the Anti-saccade task between 90ms and 1000ms after STIM appearance | 14092 | 10.22% |
| Anticipatory Correct Pro-Saccade | A correct Pro-saccade between -110ms and 89ms relative to STIM appearance | 2472 | 1.79% |
| Anticipatory Correct Anti-Saccade | A correct Anti-saccade between -110ms and 89ms relative to STIM appearance | 1236 | 0.90% |
| Anticipatory Pro-saccade Direction error | An erroneous saccade to the Anti-location during the Pro-saccade task between -110ms and 89ms relative to STIM appearance | 1914 | 1.39% |
| Anticipatory Anti-saccade Direction error | An erroneous saccade to the STIM during the Anti-saccade task between -110ms and 89ms relative to STIM appearance | 1538 | 1.12% |
| fixation break | the participant looked away from FIX prior to its disappearance and did not return prior to the end of the FIX epoch. | 7848 | 5.69% |
| no saccade | the participant fixated on the FIX but never made a saccade after STIM appearance. | 352 | 0.26% |
| random saccade | the participant fixated on the FIX and then made a saccade to a non-task relevant location after STIM appearance. | 330 | 0.24% |
| never fixated | the participant never properly fixated on the FIX | 1521 | 1.10% |
| eye loss | too much data was lost to properly analyze | 755 | 0.55% |
| **TOTAL** | Total from 592 participants (380 female) | 137896 | 100.00% |

**Figure 8:**
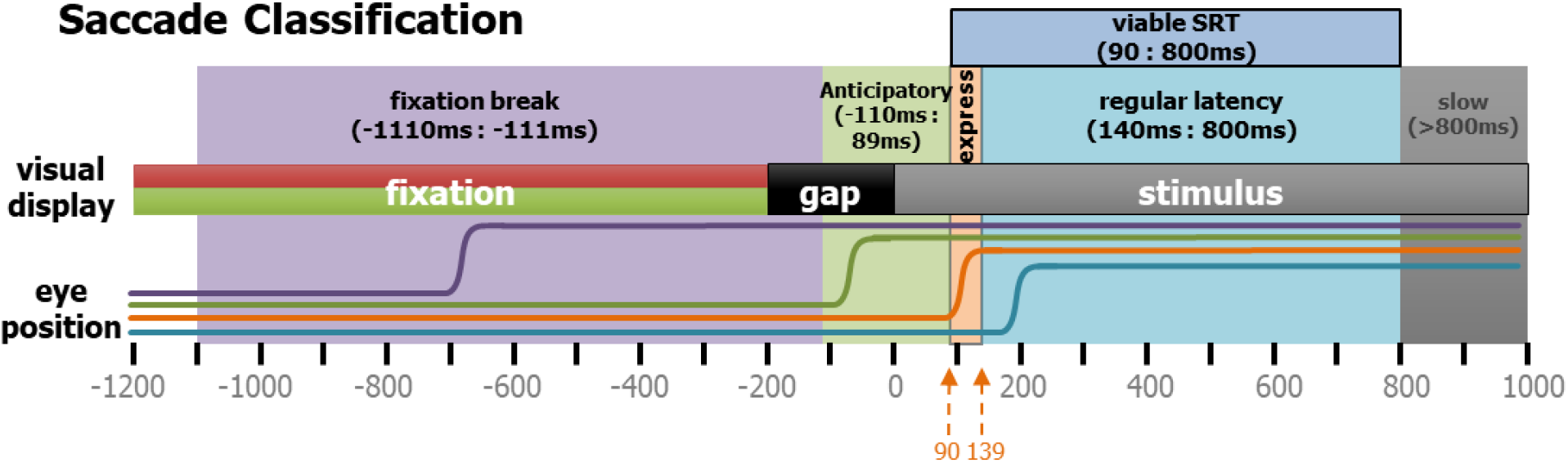
Task related Saccade Classification. Saccades in the IPAST were categorized by timing as well as direction. Here, 4 different types of rightward saccades are shown schematically. Purple indicates a fixation break, green is an anticipatory saccade, orange is an express saccade, and cyan is a regular latency saccade.

#### IPAST Trial Classification

The final goal of this analysis was to classify each trial in the IPAST. Previously, trial classification was rather rudimentary, only including correct trials, incorrect trials, and ‘other’ trials. These ‘other’ trials were not used and could include trials lost due to poor data or some other behavior not specifically defined. Here we were more thorough in defining each trial so that even the ‘other’ trials could be useful.

Because we have a larger variety of trial types, we can now calculate different behavioral metrics and produce standardized results. Table 1 contains the list of all possible trial types. Using the new categories of trial types we could calculate not only error rates, but also non-compliance rates. For example, trials that fell under no-saccade, random saccade, and never fixated, categories were summed up, and used as a measure of non-compliance. Additionally, trials with fixation breaks and anticipatory saccades could also be independently measured. Trials in which fixation briefly lapsed, i.e., fixation was broken but was re-established before the FIX turned off, were allowed to be marked by the subsequent behavior. These lapses were marked in the saccade table and could be used for further investigations. Unmarked trials and data-lost trials were completely removed and made up only 0.59% of all trials. This pipeline was able to read in data, categorize blinks, categorize saccades, automatically mark, and then categorize 99.96% of 137,896 IPAST trials from 592 participants, in approximately 2 hours (the data was on a Linux mainframe server and the code was run on a windows 10 PC with Intel^®^ Core™ i7-4770 CPU @ 3.40GHz).

We then defined ANTI ‘error rate’ as all direction error ANTI trials divided by the sum of all ANTI trials including fixation break, anticipatory, and non-compliance trials, and ‘error ratio’ as all direction error ANTI trials divided by the sum of only the correct and direction error ANTI trials). The ‘rate’ calculation asks, “of all the ANTI trials where a measurable behavior took place, how many were direction errors?”. The ‘ratio’ calculation is more similar to the calculations of the past, as it removed the fixation break, anticipatory, and non-compliance trials, and asks a slightly different question: “out of all the viable ANTI saccades (see Fig. 8) made, how often did they fail to suppress the sensory driven signals?”. For many participants these values were similar. However, for those who struggled to understand the task and have more fixation break, anticipatory, and/or non-compliant trials, the difference between these values was meaningful, and warranted further investigation. Additionally, fixation break ‘rates’, anticipatory ‘rates’, and other similar scores were calculated using all trials.

## Discussion

Here we describe in detail a new data analysis pipeline to automate and optimize data analysis from the video-based eye tracking where we can extract measures of eye position, pupil size, and blinks. These methods are not meant to be a standardized protocol for all anti-saccade research, merely a description of what we have done and why. This process is not only faster than manual marking but also far more consistent and produces repeatable measurements of behavior.

With an understanding of the nature of the PSOs, the accuracy of the spatial location saccade end-points has been improved. The described approach post-hoc combined the PSOs with the main saccade even when the speed drops below threshold in between the two events. This is done for every saccade in a task-agnostic fashion. Similarly, each and every blink is also detected and categorized. Blinks were once a nuisance to overcome in measuring eye movement via video-based tracking but have now become a behavioral marker in their own right. Additionally, the trials that were once thrown away can now be utilized. The fixation break, anticipatory, and the different non-compliance trials can be used as meaningful behavioral markers. Collectively, these steps lead to the standardization of IPAST behavior across numerous data collections sites to create easily comparable scores.

### Task Parameters

The ‘interleaved’ and ‘gap’ aspects of the IPAST were chosen for several reasons.

#### Interleaved

Previous versions of the anti-saccade task (Antoniades et al., 2013; Munoz et al., 1998) used a blocked design which isolated pro-saccades from anti-saccades in separate blocks of 100 consecutive trials or more. The block design is not ideal for contrasting pro- and anti-saccade behavior because levels of concentration, arousal, and fatigue vary between blocks of different tasks and over time in a prolonged laboratory experiment. The interleaved version of the task requires more working memory (Unsworth et al., 2004), as the current trial’s rule must be updated and remembered upon STIM presentation on a trial-by-trial basis. The dynamic aspect of the interleaved task also relieves monotony and increases trial count for healthy participants. Lastly, it also assures that the visual, spatial, and temporal aspects of the two tasks (e.g., FIX and STIM luminance and timing) were as identical as possible.

#### Gap

We also found that the ‘gap’ paradigm (200ms between FIX and STIM) improved the ability to measure the dynamic levels of inhibition needed to correctly perform the anti-saccade task (Coe & Munoz, 2017), compared to the ‘step’ (0ms between FIX and STIM) or ‘overlap’ (FIX stays on during STIM) paradigms. The presence of the gap period is known to lead to a reduction in sustained activity of fixation neurons in the superior colliculus (Dorris & Munoz, 1995; Marino et al., 2012), a reduction in reaction time known as the gap effect (Saslow, 1967), and more direction errors in the anti-saccade task (Everling et al., 1998). Additionally, only two possible target locations were used so there is a high degree of preparation, for both action and inhibition, during the task(Coe & Munoz, 2017; Dorris & Munoz, 1998) Understanding these different aspects of preparation and inhibition have improved model performance to better match human data(Coe et al., 2019).

### Post-Saccadic Oscillations

Video based eye tracking has been shown to record more PSOs than classical eye coil techniques (Kimmel et al., 2012; Frens & van der Geest, 2002). Thus, the end point of saccades as measured by video-based eye tracking can be more difficult to detect (Hooge et al., 2015; Mardanbegi et al., 2018; Nyström et al., 2013) PSOs affect the end points of saccades, and this has important implications for the accuracy of saccade metrics reported. Based on an understanding on what can happen to the eye during hard deceleration (Jacobi & Jagger, 1981), we describe a method to obtain a more accurate measure of the actual end-point for a saccade. We also went into great detail about how the eye-position data was smoothed because, similar to Mack et al. (2017, see fig 3), we have found that different smoothing techniques can actually increase the effects of the iris slosh on PSO, further complicating saccade end-point detection. Thus, when it comes to smoothing video-based eye tracking data, the simper the better.

The important idea to take away from this discussion is that to accurately measure the end position of a saccade one must wait for the oscillations to end, with the caveat that this artificially elongates the duration of saccades for individuals with greater slosh. In taking this approach, we sacrificed the accuracy of the duration measurement for better accuracy of the end position. Future work should be able to model the post-saccadic oscillations and form a better estimate of the time for the end of the saccade. Also, not unlike blinks, these PSOs caused by slosh could be another potential behavioral marker.

### Summary

In this paper we describe new automated analysis features that are important because they allow for standardized quantification of saccades, blinks, pupil response, and task performance across multiple cohorts of participants. To illustrate how important this is, it’s currently being used to analyze large, variable data sets from diverse participant cohorts (development, aging, neurodegenerative, neuropsychiatric, and long-haul COVID), different study designs (baseline, longitudinal, and clinical intervention) and from collection sites all around the world (Canada, USA, Mexico, Brazil, and Germany).

## Acknowledgements

We thank members of the Munoz labs for their comments on earlier versions of the manuscript. This work was funded by a research grant from the Canadian Institutes of Health Research (MOP-FDN-148418) and the Ontario Brain Institute. DPM was supported by the Canada Research Chair Program. Ethics protocol File No 6005163; PHGY-007-97.

## Footnotes

1 https://en.wikipedia.org/wiki/Slosh_dynamics

2 https://en.wikipedia.org/wiki/Fluid-structure_interaction

3 https://www.youtube.com/watch?v=X7ZFI2SQRyI

